# A rapid protocol for ribosome profiling of low input samples

**DOI:** 10.1101/2022.09.23.509038

**Authors:** Andreas Meindl, Markus Romberger, Gerhard Lehmann, Norbert Eichner, Leon Kleemann, Jie Wu, Johannes Danner, Maria Boesl, Mikhail Mesitov, Gunter Meister, Julian König, Sebastian Leidel, Jan Medenbach

## Abstract

Ribosome profiling provides quantitative, comprehensive, and high-resolution snapshots of cellular translation by the high-throughput sequencing of short mRNA fragments that are protected from nucleolytic digestion by ribosomes. While the overall principle is simple, the workflow of ribosome profiling experiments is complex and challenging, and typically requires large amounts of sample, limiting its broad applicability. Here, we present a new protocol for ultra-rapid ribosome profiling from low-input samples. It features a robust strategy for sequencing library preparation within one day that employs solid phase purification of reaction intermediates, allowing to reduce the input to as little as 0.1 pmol of RNA. Hence, it is particularly suited for the analyses of small samples or targeted ribosome profiling. Its high sensitivity and its ease of implementation will foster the generation of higher quality data from small samples, which opens new opportunities in applying ribosome profiling.

## Introduction

Ribosome profiling generates genome-wide snapshots of cellular translation based on the analysis of short mRNA fragments that are protected from nucleolytic digestion by ribosomes. These ribosome footprints or ribosome-protected fragments (RPFs) yield the precise position of translating ribosomes on mRNAs at the moment of cell lysis (Steitz 1969) and their highly parallel sequencing yields comprehensive and quantitative insight into translation dynamics (Ingolia et al. 2009).

Ribosome profiling has significantly advanced our understanding of the plasticity and regulation of translation during development or in response to stimulation. It has provided evidence for pervasive translation of RNAs and revealed translation outside previously annotated protein-coding regions, thus spurring the discovery of novel peptides. Moreover, its high resolution has unveiled codon-dependent differences in elongation rates and ribosome collisions (Ingolia 2014; Nedialkova and Leidel 2015; Ingolia et al. 2019) and selective ribosome profiling has provided deep and novel insights into the mechanism of translation (Andreev et al. 2021). For the latter, select sub-populations of ribosomes are purified prior to the analyses of the associated footprints; these include e.g. initiating ribosomes (Archer et al. 2016; Bohlen et al. 2020; Wagner et al. 2020) or ribosomes associated with specific proteins like cotranslational chaperones (Oh et al. 2011; Doring et al. 2017).

Numerous variations have been introduced to the protocol, adapting it to a wide range of model organisms and tailoring it to address different aspects of translation and its regulation. This includes the use of different nucleases (Gerashchenko and Gladyshev 2017) or translation inhibitors to stall ribosomes on initiation codons (Harringtonine or Lactimidomyin) or at specific stages during the translation cycle (e.g. Cycloheximide, Anisomycin, Emetine, Chloramphenicol or Tigecyclin) (Ingolia et al. 2011; Lee et al. 2012; Wu et al. 2019). Furthermore, different methodological approaches have been employed for the preparation of ribosomal monosomes that contain ribosome-protected fragments. This includes enrichment of monosomes by sucrose density gradient centrifugation (Ingolia 2010), via sucrose cushions (Lacsina et al. 2011; Ingolia et al. 2012), or by size exclusion chromatography using spin columns (ARTSeq protocol, Illumina), but also the affinity purification of elongating ribosomes via puromycin derivatives that are integrated into the nascent peptide chain (Clamer et al. 2018).

Even though the original ribosome profiling protocol produces datasets of excellent quality, it is complex and challenging to perform. A careful fine-tuning of the nucleolytic digestion, the purification of ribosomal complexes, and the precise size selection of RNA for library preparation are critical for the generation of high-quality datasets. To date, numerous ribosome profiling protocols exist that vary in numerous experimental details, and, as many of the experimental steps have not been standardized, a comparison of the data generated by the individual workflows is difficult.

Moreover, in the case of low-input samples, the efficiency of library preparation is often becoming limiting. This is particularly the case for selective ribosome profiling where specific ribosomal subpopulations are analyzed. Similarly, small tissue samples (e.g. from clinical biopsies) have proven difficult to profile. To overcome these challenges, a standardized ribosome profiling protocol with a sensitive and efficient sequencing library preparation is highly desirable.

Here we present a novel library preparation protocol for ribosome profiling that allows the robust generation of high-quality libraries from as little as 0.1 pmol of RPFs. The simplified workflow produces high-quality libraries at little effort and within 24 h (Figure 1). This is achieved by combining the enzymatic steps of the original protocol with recently introduced innovations from different sequencing library preparation protocols into a streamlined and optimized workflow: (1) solid-phase reversible immobilization (SPRI) offers convenient and rapid purification of reaction intermediates while minimizing loss of material (2) an optimized strategy for PCR amplification prevents overamplification to ensure the high quality of the final libraries (3) highly efficient ligation of adapters and newly designed primers enhance the sensitivity and facilitate experimental multiplexing (Figure 2) (4) degenerate nucleotides in the ligation adapters reduce ligation bias and allow the identification of PCR duplicates during data analysis (Lecanda et al. 2016). In summary, the presented protocol is simple to perform, cost efficient, more sensitive, and faster than the original protocol.

**Figure 1:**
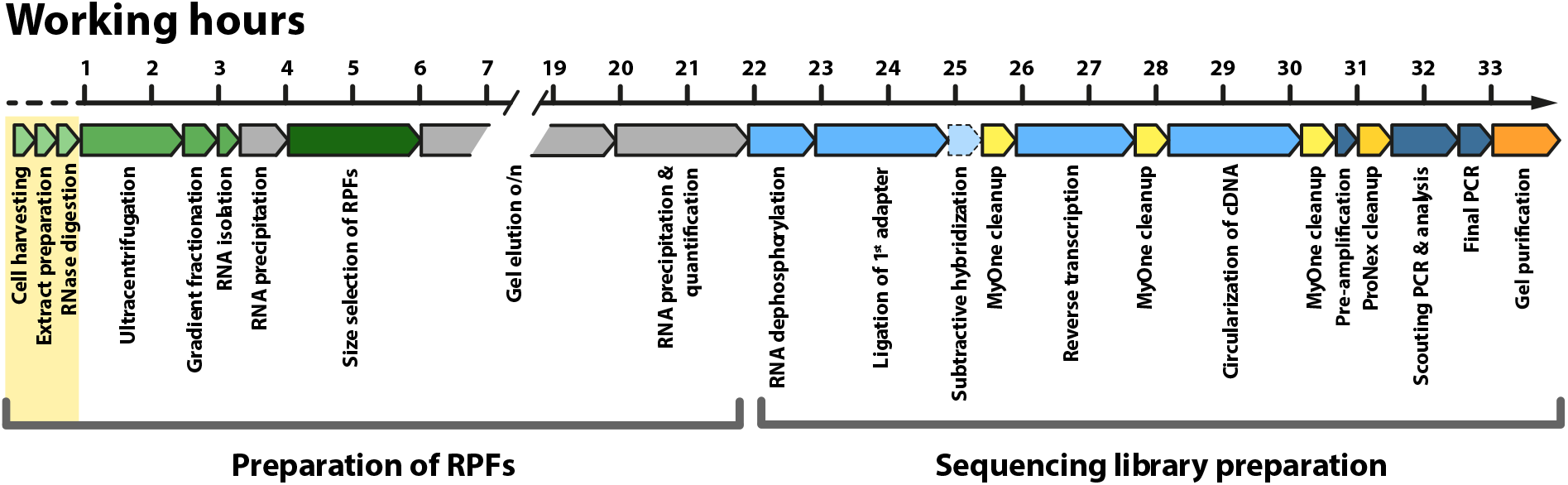
Overview of the workflow. As in the original protocol, ribosome protected fragments (RPFs) are produced by nucleolytic treatment of native cell extracts and purified by sucrose gradient density centrifugation and size selection. For sequencing library preparation, terminal phosphates are enzymatically removed from the RNA, followed by ligation of an adapter to the 3’ end to support reverse transcription. After circularization of the cDNA, the product is amplified by PCR to generate an amplicon for Next Generation Sequencing on an Illumina platform. Purification of reaction intermediates occurs mostly via solid phase extraction with magnetic beads. Optionally, a depletion of rRNA-derived, contaminating sequences by subtractive hybridization can be integrated at a strategically optimized position into the workflow (box with dashed outline). To adapt the protocol to different samples, only the initial steps require optimization (highlighted in yellow). Working hours are given at the top.

## Methods

### Recombinant protein production

The open reading frame encoding TS2126 Ligase with a C-terminal His-tag was synthesized [Geneart] and cloned into pET22B using the NdeI and EcoRI restriction sites. Expression and purification occurred according to Blondal et al. 2005. To produce T4 RNA ligase 2 the sequence encoding amino acids 1-294 was amplified by PCR from *Escherichia coli* and cloned into pET16B using the NdeI and BamHI restriction sites. Subsequently, the point mutations R55K and R227Q were introduced by PCR mutagenesis. Expression and purification followed the protocol by Ho et al. 2004.

### Preparation of ribosome-protected fragments

#### *H. sapiens* K562 cells

K562 cells were grown in DMEM supplemented with 10 % FCS at 37°C and 5% CO_2_ to a maximal density of 1-1.25 million cells per ml. K562 cells were treated for 3 min with 100 μg/ml Cycloheximide and subsequently harvested by centrifugation for 2 min at 2,000 × g and 4°C. The cell pellet was resuspended in 300 μl lysis buffer (5 mM Tris/Cl pH7.4, 1.5 mM KCl, 5 mM MgCl_2_, 0.5% Triton X-100, 0.5% Sodiumdeoxycholate, 1 mM DTT, 100 μg/μl Cycloheximide, 10 U/ml Turbo DNase [ThermoFisher], 100 μM RNaseOUT [ThermoFisher], 1 × cOmplete protease inhibitor cocktail [Roche]) and incubated on ice for 15 min. After centrifugation at 14,000 × g for 10 min at 4°C, the cell lysate was transferred into a new reaction tube and UV absorption was determined at 260 nm in a spectrophotometer. In parallel, the RNA concentration of the lysate was determined using the Qubit RNA BR Assay-Kit and a Qubit fluorometer [ThermoFisher]. For nucleolytic digestion, 8 U of RNase I [ThermoFisher] were used per AU260 of lysate and digestion was performed for 15 min at 20°C under slow rotation. The reaction was stopped by the addition of 100 U of SUPERase•In RNase Inhibitor [ThermoFisher].

#### Density gradient ultracentrifugation

After nucleolytic digestion, the samples were subjected to density gradient ultracentrifugation on 10-50% linear sucrose gradients in a buffer containing 20 mM Tris/Cl pH7.4, 150 mM NaCl, 5 mM MgCl_2_, 1mM DTT, and 100 μg/ml Cycloheximide. Centrifugation was as follows: SW40Ti rotor: 3 h at 35,000 rpm (155k × g) at 4°C, SW55Ti rotor: 1.5 h at 50,000 rpm (237k × g) at 4°C, SW60Ti rotor: 70 min at 50,000 rpm (257k × g) at 4°C [all rotors from Beckmann]. The gradients were fractionated into 1 ml (SW40 rotor) or 400 μl fractions (SW55 and SW60 rotors), under continuous UV monitoring using a siFractor fractionation unit [siTOOLs Biotech] connected to an ÄKTApurifier FPLC [Cytiva].

For baseline correction, a sliding window approach was employed. From a reference point, starting at the beginning of the elution profile, it searches for the lowest absorption in a defined elution volume (1.5 ml) downstream. If all absorption values within the window are higher than the reference point, the lowest value is used as the next reference point and calculation is repeated at this point. If absorption values lower than the reference point are detected within the search window, the next point for calculation is the one at the steepest angle below the reference point. Finally, all calculated reference points are connected to generate a profile that is subtracted from the data.

#### Purification of RPFs

The fraction containing the 80S monosomes was diluted with an equal volume of nuclease-free water end extracted with phenol:chloroform:isoamylalchol (25:24:1). The aqueous phase was transferred to a new reaction tube and RNA was precipitated with 2.5 volumes ethanol in the presence of 3 μl of linear acrylamide solution [ThermoFisher]. Subsequently, the RNA was separated by denaturing PAGE (15%) and visualized by SYBR Gold [ThermoFisher] staining. RNAs in a size range from 26 to 30 nts were excised from the gel and eluted in a buffer containing 300 mM NaOAc, 1 mM EDTA, 0.25% SDS, and 20 U/ml SUPERase•In RNase Inhibitor. After precipitation, the RNA was resuspended in nuclease free water and the concentration was determined using a Qubit Fluorimeter with the microRNA Assay Kit [ThermoFisher].

#### *S. cerevisiae* BY4247

BY4742 yeast cells were grown at 30°C in YPD medium. Whole cell extracts were prepared from exponentially growing cultures essentially as described in Schwank et al. 2022. In brief, at an OD600 of 0.6-0.7, the cells were harvested by centrifugation and flash frozen in liquid nitrogen. Cell lysis occurred mechanically under cooling with liquid nitrogen (Cryolys Cooling System) using 0.75-1mm glass beads in a Precellys tissue homogenizer at 6,000 rpm in a buffer containing 20 mM Tris/Cl pH 7.4, 150 mM NaCl, 5 mM MgCl_2_ 1% Triton-X100, 1 mM DTT, 25 U/ml Turbo DNase, and 100 μg/ml Cycloheximide. After clearing the extract by centrifugation at 20,000 × g for 10 min at 4 °C, it was subjected to RNase I digestion for 5 min on ice using 30U of enzyme per AU_260_.

Subsequently, ribosomal complexes were separated by sucrose density ultracentrifugation as described in Pospisek and Valasek 2013. In brief, 15-50% linear sucrose gradients were prepared in a buffer containing 20 mM Tris/Cl pH 7.4, 150 mM NaCl, 5 mM MgCl_2_, 1 mM DTT, and 100 μg/ml Cycloheximide. Ribosomal complexes were separated by centrifugation in a SW40Ti rotor for 2.5 h at 35,000 rpm at 4°C or in a SW55 rotor for 1.5h at 50,000 rpm at 4°C. Fractionation of the gradients and purification of the ribosomal footprints occurred as described above.

### Sequencing library preparation

#### Optimized protocol

The indicated amounts of gel-purified RNA were dephosphorylated by treatment with 5 U of T4 PNK [New England Biolabs] in a 20 μl reaction supplemented with 1x reaction buffer and 20 U of RNaseOUT [ThermoFisher] for 30 min at 37°C. After heat inactivation of the enzyme, the reaction was supplemented with 4.5 μl DMSO, 6 μl 50% PEG400, 1.5 μl RNA-ligase buffer, 1 μl T4 RNA Ligase 2 (aa 1-294, R55K, K227Q) and 20 pmol of rApp-L7 linker oligonucleotide. The reaction was incubated over night at 16°C followed by an incubation for 1h at 37°C and by heat inactivation of the enzyme. For depletion of contaminating rRNA-derived sequences, the entire reaction was subjected to subtractive hybridization using a riboPOOL kit [siTOOLs Biotech] according to the manufacturer’s instructions. Subsequently, the ligation product was extracted using 20 μl MyONE Silane beads [ThermoFisher] in 650 μl RLT Buffer [Qiagen] and 720 μl ethanol. Reverse transcription was performed as described in Buchbender et al. 2020, using 0.5 pmol P7 RT oligonucleotide. For hydrolysis of the RNA, the reaction was supplemented with 1.65 μl of 1M NaOH and incubated for 20 min at 90°C, followed by neutralization with 20 μl of 1M HEPES/KOH pH7.3. The cDNA was purified with 10 μl of MyONE Silane beads as described above and eluted in 14 μl of nuclease-free water. Circularization of the cDNA occurred in a 20 μl reaction containing 0.5 mM ATP, 2.5 mM MnCl_2_, 0.05 mM ATP, 50 mM MOPS (pH 7.5), 10 mM KCl, 1 mM DTT, and 1μl TS2126 ligase for 2h at 60°C followed by an incubation for 10 min at 80°C. The circular product was extracted with 10 μl MyOne Silane beads in 650 μl buffer RLT and 720 μl ethanol. After elution it was subjected to a pre amplification by PCR using the P5_s and P7_s primers and KAPA HiFi HotStart ReadyMix [Roche] according to the manufacturer’s instructions. The PCR product was purified using the ProNex size-selective purification system [Promega] with a 1:2.95 v/v ratio of sample to beads. Scouting PCRs and final amplification of the library were performed as described in Buchbender et al. 2020. In brief, for scouting PCRs, 10 μl reactions were set up using the KAPA HiFi HotStart ReadyMix [Roche] with P5 and TrueSeq P7 indexed primers and subjected to 6-16 cycles of PCR. After analysis of the PCR products either by native PAGE or a high sensitivity screen tape assay on a TapeStation [Agilent], optimal PCR conditions were chosen according to the concentration of the PCR product and the appearance of daisy chains (compare Huppertz et al. 2014). Final amplification of the library occurred in a 40 μl reaction using the conditions determined in the scouting runs. Subsequently, for the removal of primers and the depletion of empty amplicons, the PCR reaction was subjected to purification using a Blue Pippin [Sage Science] with 3% agarose cassettes selecting fragments of a length of 168bp with the setting ‘tight’. All primer sequences are listed in Supplemental Table 1.

Comparison with the ‘classical’ library preparation protocol: HEK293 cells were harvested as described in Kim et al. 2021 with the following adaption: the lysis buffer was supplemented with 25U/ml Turbo DNase [ThermoFisher] before use. Ten A260 units of lysate were digested with 1000U RNase I [ThermoFisher]. After RNA dephosphorylation by PNK, the samples were split into two tubes to perform the Kim et al. protocol with one half and this new protocol with the other half. Ribosomal RNA was depleted using biotinylated oligonucleotided as described before (Ingolia et al. 2012), Supp. Tab. 2.

#### iCLIP2 protocol

RPFs were subjected to library preparation as described (Buchbender et al. 2020) with the following modification: ligation of the first adapter occurred in solution in a 20 μl reaction containing 1 × T4 RNA ligase buffer [New England Biolabs], 20 U RnaseOUT [Invitrogen], 10 U T4 RNA ligase 10 [New England Biolabs], 4 μl PEG400 [Sigma Aldrich]. Subsequently, the ligation product was purified via MyONE Silane beads and the protocol was continued as described in Buchbender et al. 2020.

### Sequencing and data analyses

For sequencing, libraries were quality controlled using a high sensitivity screen tape assay on a TapeStation [Agilent] and quantified using the KAPA Library Quant Kit [Roche]. Sequencing occurred on a MiSeq (V3 kit, 150 cycles, 90 cycles single end, 6 nt index read).

For bioinformatic analyses, R version 3.6.3 was used. After de-multiplexing of the sequencing reads, adapter trimming occurred with Cutadapt (version 2.8, parameters: adapter=AGATCGGAAGAGCACACGTCT, overlap=10, minimum-length=0, discard-untrimmed). UMI tools (version 1.0.1) was used to extract the UMIs and to associate their sequence with the read identifier. Trimmed reads were mapped against hg38 (*H. sapiens*) or R64 (sacCer3, *S. cerevisiae*) using bowtie2 (version 2.4.1) using the standard parameters. PCR duplicates were subsequently identified using UMI tools (parameters: --extract-umi-method read_id --method unique).

Metagene plots at initiation and termination codons and periodicity analyses of RPFs were performed as described in Sharma et al. 2021.

## Results

### Efficient sequencing library preparation from minute amounts of RNA

We have developed an improved and rapid strategy for the generation of sequencing libraries from minute amounts (0.1 pmol) of ribosome-protected fragments (Figure 1). The original ribosome profiling protocol employs polyacrylamide gel electrophoresis (PAGE) for the purification of reaction intermediates which is laborious and time consuming (Ingolia et al. 2009; Ingolia 2010; Ingolia et al. 2012). Furthermore, elution of the sample from gel slices typically results in significant sample loss, particularly when working with small amounts of RNA (we typically recover ≤30% of a sample of 30 pmol of RNA, data not shown). As an alternative to gel purification, solid-phase reversible immobilization (SPRI) has been introduced, e.g. for sequencing library preparation from samples obtained from cross-linking immunoprecipitation experiments (iCLIP2, Buchbender et al. 2020)). Extraction with Dynabeads MyOne silane is rapid, robust, and yields a higher sample recovery in comparison to gel purification.

Building on the iCLIP2 library preparation approach, we developed a protocol tailored specifically to the sequencing of small RNAs derived e.g. from ribosome profiling experiments. Besides employing solid-phase extraction of reaction intermediates, we re-designed the adapter sequences, allowing the generation of shorter amplicons with P5 and P7 adapters. These amplicons are similar to the ones produced by the original ribosome profiling protocol but lack several nucleotides adjacent to the insert that are not strictly required for sequencing (Figure 2). In contrast to iCLIP2-derived libraries, for experimental multiplexing, in-line barcoding is replaced by a P7-side encoded index that is introduced during the final PCR. This reduces primer costs and facilitates multiplexing with libraries derived from different experimental approaches such as e.g. RNA sequencing for expression profiling.

**Figure 2:**
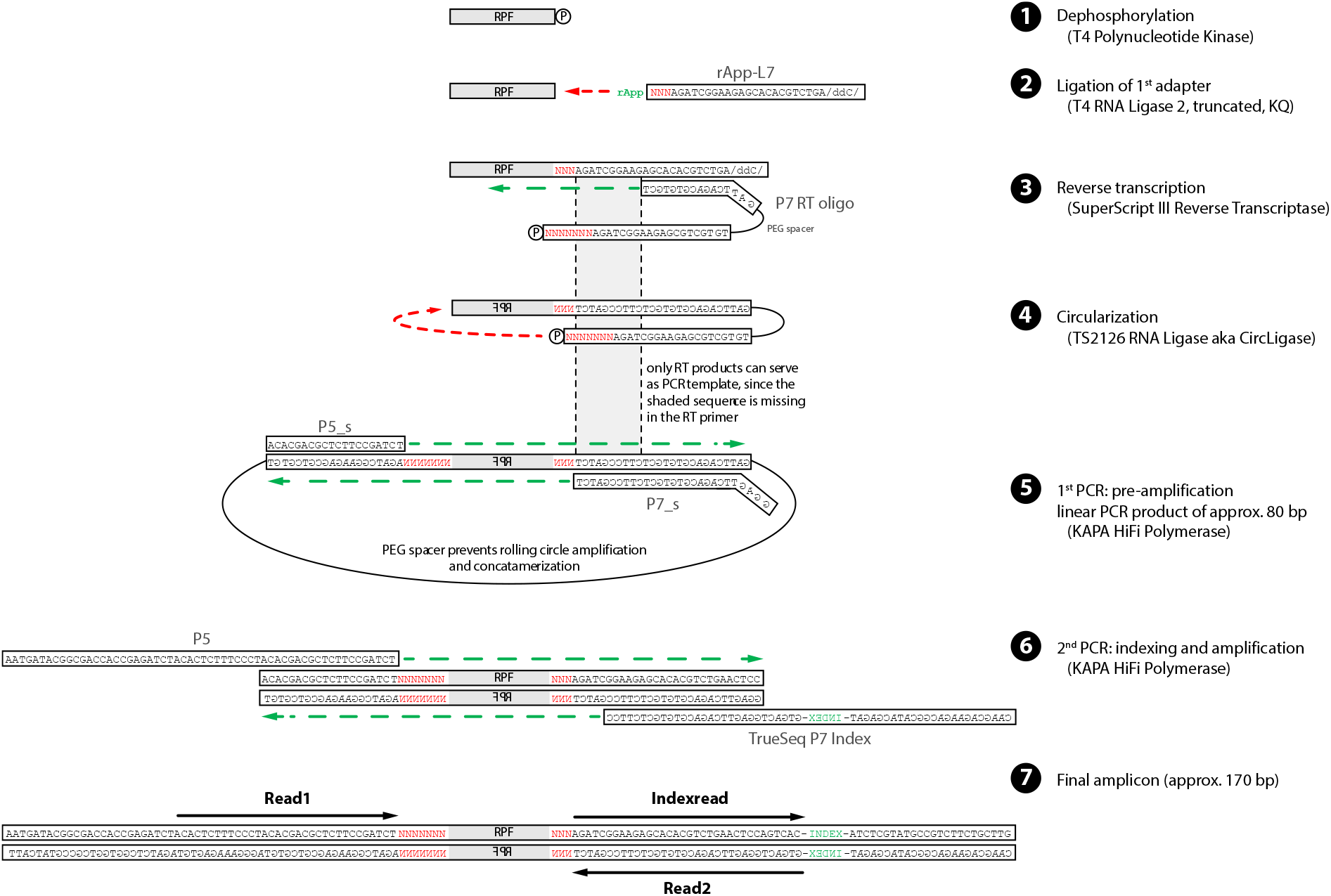
Schematic overview of sequencing library preparation. Individual steps in the protocol are described on the right; identity of the enzymes employed in each step is provided in brackets. Degenerate nucleotides that reduce ligation bias and that are used for the identification of PCR duplicates are highlighted in red, the position of an index for experimental multiplexing in green.

In agreement with previous reports, ligation of the first linker to the RNA fragments by T4 RNA ligase I has proven inefficient in our hands (Suppl. Figure 1A) (Ho et al. 2004; Pfeffer et al. 2005). We therefore opted for T4 RNA ligase II (truncated aa 1-294 and carrying the mutations R55K and R227Q). In the presence of PEG400 and DMSO, ligation of 5 pmol of a 30 nt RNA oligonucleotide to a preriboadenylated adapter proceeds almost to completion (Suppl. Figure 1A). Moreover, as in the original ribosome profiling protocol, we employ circularization of the cDNA instead of ligation of a second adapter. The efficiency of circularization with TS2126 ligase (aka CircLigase) is effective and comparable in efficiency to ligation of a second adapter with T4 RNA ligase I and avoids known ligation biases (Suppl. Figure 1B & C) (Lecanda et al. 2016). Furthermore, the reaction proceeds much faster, requiring only a 2 h incubation instead of an overnight reaction and the enzyme TS2126 works at elevated temperatures, which reduces the impact of local secondary structures on ligation and it also exhibits no significant preference in the positions surrounding the ligation site, reducing bias during sequencing library generation (Lama et al. 2019). To further minimize biases, the ligation adapters contain degenerate positions at their 5’ ends (Figure 2) (Kim et al. 2019). Simultaneously, these sequences serve as unique molecular identifiers (UMIs) for the identification of PCR duplicates during data analysis after sequencing.

To evaluate the performance of the optimized library preparation protocol, we compared it to the recently developed iCLIP2 protocol. We reasoned that when using small amounts of input material for library preparation, an inefficient strategy will necessitate extensive PCR amplification to yield sufficient material for sequencing. This results in an increase of the fraction of PCR duplicate reads that can be identified during bioinformatic analysis based on mapping position and unique molecular identifier (Suppl Figure 2A). In contrast, a library generated by an efficient protocol should contain very few PCR duplicates. Legacy sequencing library preparation protocols employed for ribosome profiling typically require 10-20pmol of RNA input for production of a high-quality library (Ingolia et al. 2009; Ingolia 2010; Ingolia et al. 2012). We therefore subjected 5 pmol of purified RNA (26-30 nt in length, generated by RNAse I digestion) to sequencing library preparation using either the iCLIP2 protocol or our optimized library preparation protocol. The libraries were amplified to yield similar amounts of product and sequenced at comparable depth. Analysis of the sequencing data revealed that the iCLIP2-derived libraries are dominated by PCR duplicates (~70%), while the optimized protocol produces high-quality libraries that contain less than 2% PCR duplicates (Figure 3A).

**Figure 3:**
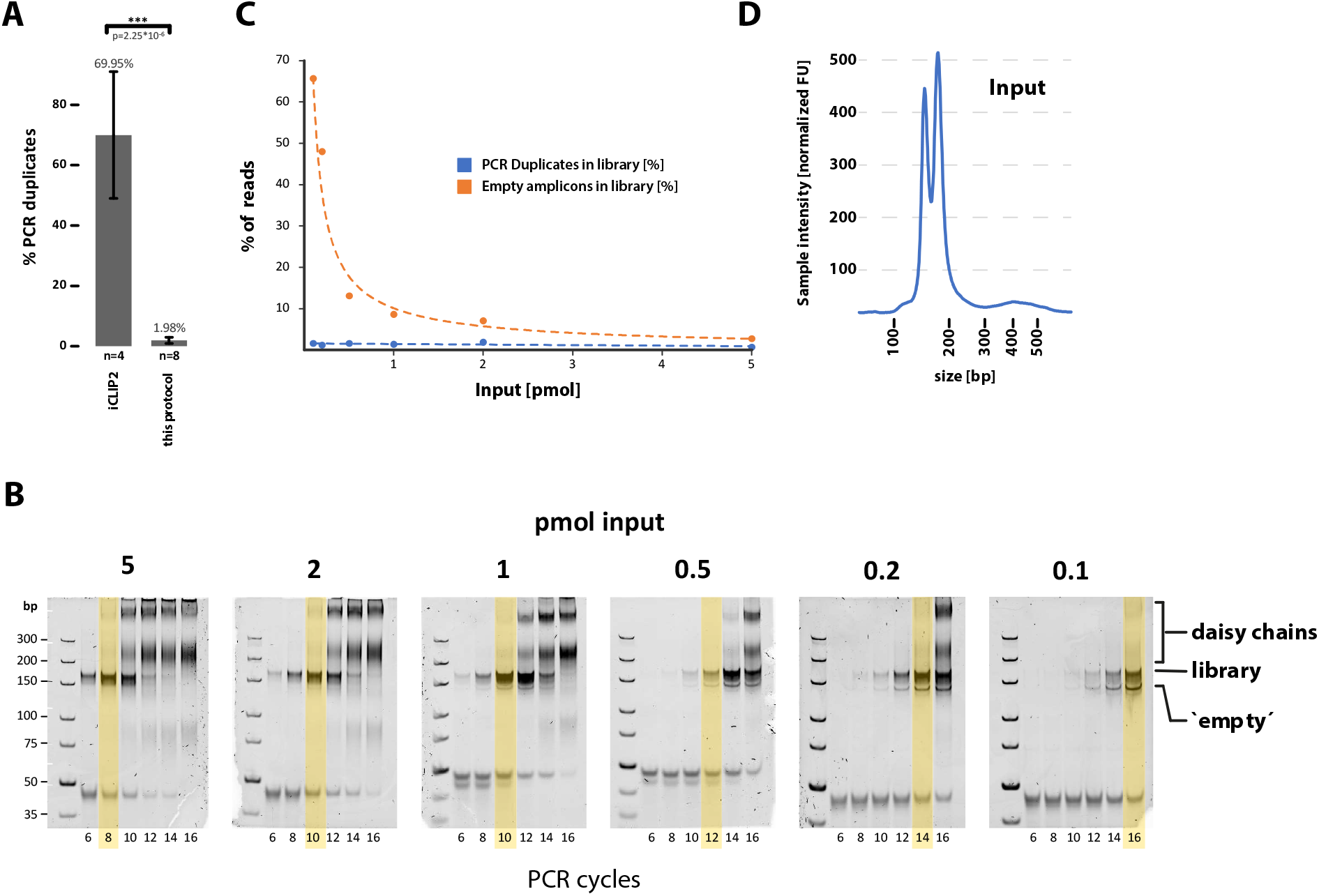
Sequencing library preparation from minute amounts of sample. (A) Analysis of PCR duplicates from sequencing libraries generated from 5 pmol of RNA (26-30 nt in length, generated by RNase I digestion) using the iCLIP2 library preparation protocol (left), or the protocol described here. Number of replicates are provided below each bar, depicted are mean +/− SD. (B) Results of scouting PCRs of libraries prepared from different amounts of input material (as noted above each gel). Provided below each lane are the number of PCR cycles that were performed. The positions of high molecular weight heteroduplex DNA (‘daisy chains’, derived from over-amplification of the libraries) and ‘empty’ amplicons are indicated on the right. Yellow boxes indicate conditions that have been used to generate the final samples for high throughput sequencing. (C) Analysis of PCR duplicates and ‘empty’ amplicons contained in the libraries depicted in panel B. (D) Depletion of ‘empty’ amplicons by automated agarose gel electrophoresis. Sample prior (left) and after purification (right).

We next tested the sensitivity of the optimized library preparation protocol by subjecting different amounts of purified ribosome-protected fragments (ranging from 5 to 0.1 pmol) to library preparation. In all cases we obtained sequencing libraries and, as expected, low amounts of input material require an increased number of PCR cycles (Figure 3B). After sequencing, analyses of the PCR duplicates in the libraries revealed that even with minute amounts of input material (0.1 pmol), duplicate reads do not exceed 2% (Figure 3C). This demonstrates that the protocol is well suited for the generation of sequencing libraries from as little as 0.1 pmol of RNA (corresponding to approx. 1ng of 30 nt RNA).

With decreasing amounts of input material, we detect an exponential increase in amplicons that lack an insert. These ‘empty’ amplicons are generated from adapters (rApp-L7) that fail to ligate to an RNA molecule and which provide a template for the extension of the RT oligonucleotide (P7 RT oligo) during reverse transcription (Figure 2 and Suppl. Figure 2B). After circularization, non-extended RT oligonucleotides are not subject to PCR amplification as the PCR primer targets a sequence that is generated during reverse transcription. In contrast, the partially extended RT oligos efficiently engage in PCR amplification, resulting in the generation of ‘empty’ amplicons. Different strategies can be employed for the removal of these unwanted sequences. (1) In the first ligation reaction, the concentration of the rApp-L7 adapter can be adjusted to the amount of input material. A decrease in the concentration of the adapter oligonucleotide, however, can significantly reduce ligation efficiency. (2) Non-ligated adapters can be degraded enzymatically using a 5’ deadenylase and the DNA-specific exonuclease RecJ. While this reaction works efficiently, we reasoned that removal of the adapters at an early stage will result in an increased loss of sample during the workflow due to unspecific adsorption onto the plasticware (the leftover adapters can block surfaces, reducing adsorption of the actual sample). For removal of the empty amplicons, we therefore introduced a final purification step using an automated agarose gel electrophoresis system (Blue Pippin, Sage Science). It allows the efficient depletion of empty amplicons prior to sequencing (Figure 3D & Suppl Figure 3), decreasing their abundance from ~70% (0.1 pmol input) to approx. 1 % (N=4) in the final sequencing library.

To test whether this library preparation strategy can be easily adopted by additional labs, samples were processed in a non-developer lab. The results after library preparation were comparable between both labs (Figure 3 and Suppl Figure 4) attesting to the simple implementation of the protocol.

The final data exhibit features of high-quality ribosome profiling datasets. In metagene analyses, data generated by the new protocol is indistinguishable from data that was obtained from the same sample using the classical library preparation strategy. Reads are strongly enriched in open reading frames compared to untranslated regions with characteristic peaks at translation initiation and -termination codons (Figure 4A). Moreover, depending on the degree of nucleolytic digestion, a characteristic three-base periodicity for RNA fragments of 27, 28, and 29 nt in length can be observed (Figure 4B), which is characteristic for RPFs (Ingolia et al. 2009).

**Figure 4:**
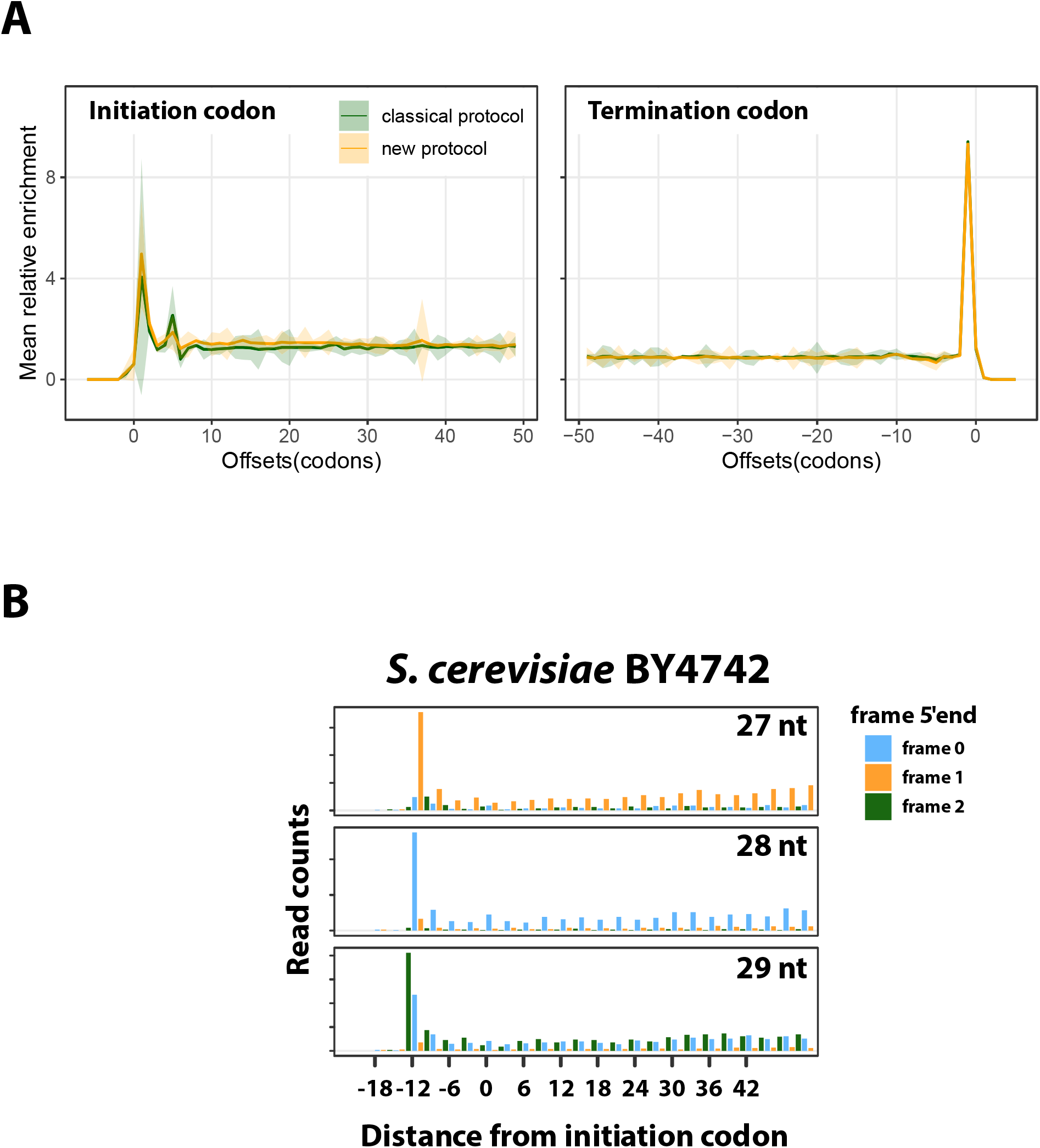
The novel protocol generates high quality ribosome profiling data. (A) Metagene analyses of read depth around translation initiation (left panel) and termination codons (right panel). Compared are libraries generated with the classical protocol (Ingolia 2010; Kim et al. 2021) (shown in green) versus libraries generated with the novel protocol (shown in yellow). Depicted are mean values (solid line) ± 95% confidence interval (shaded area). (B) The 5’ ends of ribosome footprints generated with this protocol from *S. cerevisiae* exhibit 3 base periodicity. Footprint lengths of 27, 28, or 29 nt in length (as indicated in each panel) were separately analyzed and read counts for each position relative to the initiation codon are plotted. For simplicity each frame is color coded as shown on the right.

### Ribosome profiling from low input samples

The high sensitivity of the newly developed library preparation protocol allowed us to reduce the amount of input material used for the preparation of ribosome-protected fragments. For purification of ribosomal complexes after nucleolytic digestion, we routinely employ sucrose density gradient ultracentrifugation. This allows to simultaneously assess the quality of the sample, the degree of the nucleolytic digestion and the yield. Using continuous UV monitoring during fractionation of the density gradients, we could separate and harvest ribosomal complexes from small amounts of cellular extract (0.625 AU_260_ containing approx. 15 μg of total RNA as measured by fluorometry) with high accuracy and reproducibility (Figure 5A, Suppl Figure 5).

**Figure 5:**
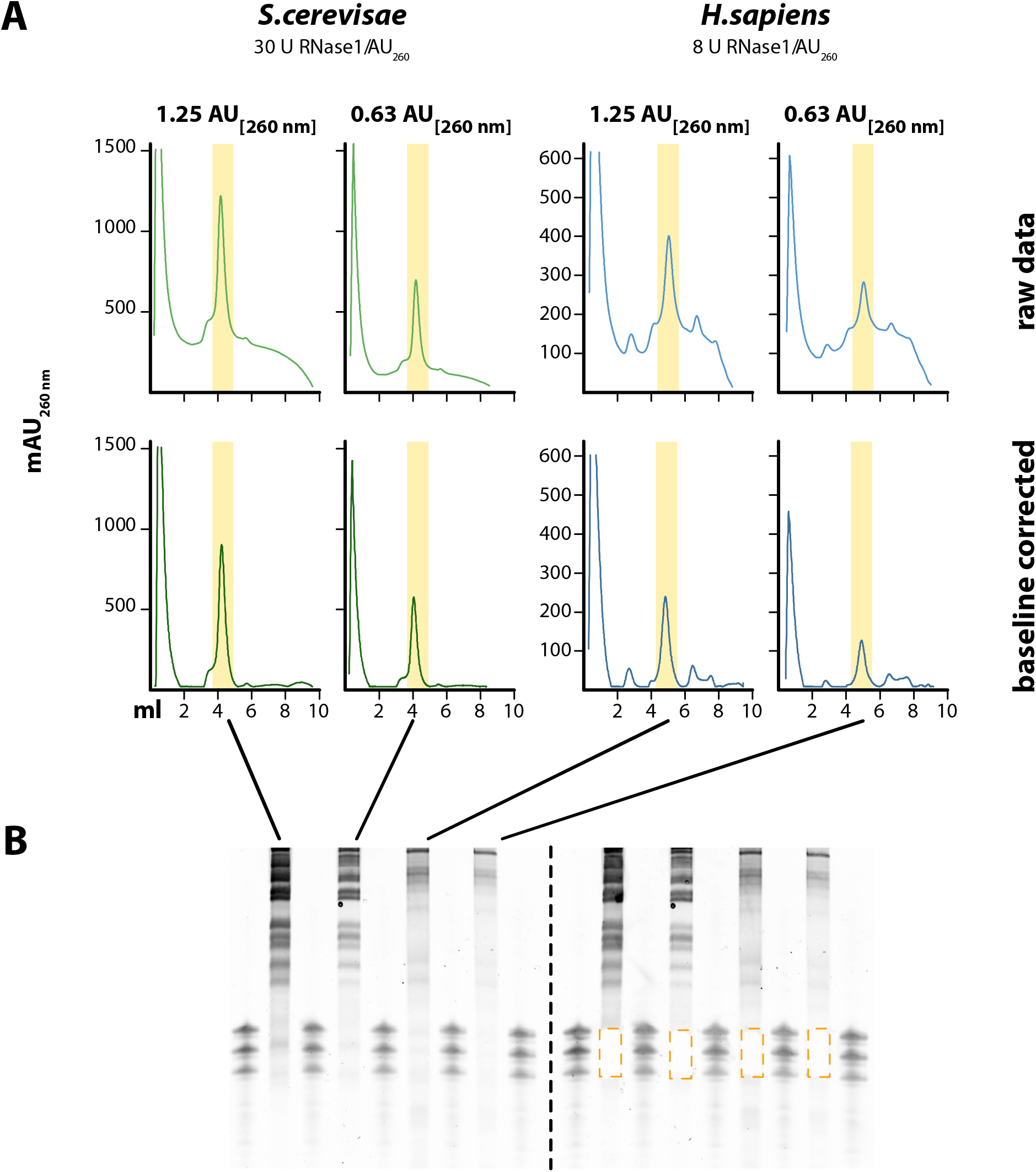
Preparation of RPFs from minute amounts of sample. (A) Sucrose density gradient ultracentrifugation of nuclease digested extracts from S. *cerevisiae* (BY4742) and *H. sapiens* K562 cells. Different amounts of extracts (1.25 AU and 0.63 AU corresponding to ~30 and ~15 μg of RNA) were separated on linear 10-15 % sucrose gradients and fractionated while monitoring UV absorption (λ = 260 nm). Raw (top) and baseline-corrected (bottom) UV profiles are depicted. Fractions containing 80S ribosomal monosomes (highlighted in yellow) were subjected to denaturing PAGE (B) to purify RNA fragments of 26-30 nt in length. Gel prior to (left) and after excision (right) of the RNA fragments.

With reduction of the input material, we observed a gradual distortion of the UV profile which is caused by differences in UV absorption through the gradient, the buffer used for cell lysis, and components of the cell lysate other than RNA. To enable a better comparability of the gradient profiles, we therefore introduced a baseline correction. Despite the inherent limitations of the mathematical correction, a better visualization of the data, such as the precise positions of the peaks, can be achieved. It performs reasonably well as determined by the quantification of ribosomal monosomes from a serial dilution (Suppl. Fig 5).

To enable accurate size selection of ribosome protected fragments from denaturing acrylamide gels, we furthermore developed a molecular weight marker. It guides the precise selection of RNA fragments between 26 to 30 nucleotides in length for preparation of the sequencing library, even if by eye no sample is visible in the 30 nt region of the gel (Figure 5B). Also, protective groups have been introduced that prevent ligation of the molecular weight marker molecules. Hence, a potential carryover of the marker does not contribute to the generation of amplicons during sequencing library preparation.

## Discussion

Ribosome profiling experiments are tailored to address various research questions and they are typically optimized for different model organisms and sample types. Parameters that vary between experiments typically include the application of translation inhibitors, different protocols for sample harvesting or preparation of native extracts, and the conditions and the choice of the enzyme for limited nucleolytic digestion to generate the ribosome footprints. All subsequent steps, however, can in principle be standardized.

Here we provide such a standardized protocol for ribosome profiling starting from purification of ribosome protected fragments after nucleolytic digestion to the production of the final sequencing library. It is extremely rapid, cheap, broadly applicable and performs robustly even with minute amounts of input. Density gradient ultracentrifugation and scouting of PCR conditions allow quality control after critical steps to ensure processing of only high-quality samples. Due to its high reproducibility, sequencing libraries produced by this protocol are highly comparable which facilitates comparisons between different experiments, which has previously been hampered by the large experimental variation between individual protocols.

High sensitivity and precision during preparation of ribosomal monosomes and accurate size selection of RNA fragments contribute to the quality of the experiment and limit contaminations derived from ribosomal RNA. However, both steps become challenging when performing ribosome profiling with low input samples that exhibit little UV absorbance. To meet these challenges, we have developed a gradient fractionation device that can be integrated into an FPLC system, enabling detection of UV absorption with high sensitivity and precise fractionation of the sample into small volumes.

Furthermore, an improved molecular size marker for gel electrophoresis guides the precise excision of low input RNA fragments with sizes of 26-30 nucleotides that are hardly visible on the gel by eye (even after staining with sensitive dyes such as SYBR-Gold). Protection groups prevent marker-derived nucleotides from contributing to the sequencing library, suppressing contaminations.

We typically perform ribosome profiling experiments using cell lysates that contain as little 0.625 AU260 (approx. 15 μg of total RNA as determined by fluorometry). Dependent on the degree of nucleolytic digestion, this yields 0.7 – 1 pmol of ~30 nt RNA fragments for library preparation. Since our improved library preparation protocol performs well with an input as little as 0.1 pmol of RNA fragments, future advances will enable the processing of even smaller samples. Our protocol will therefore stimulate analyses of translation from samples that previously were not amenable to this type of methodology - e.g. FACS-sorted cells or patient derived samples such as small tissue biopsies, organoids, or cells - as well as low input applications such as selective ribosome profiling that targets sub-populations of ribosomes.

## Data availability

All sequencing data have been deposited at SRA and are accessible via the following BioProject: PRJNA883055.

## Acknowledgements

We thank Joachim Griesenbeck for support with the experiments using S. *cerevisiae*.

