## Supplemental Information for "A rapid protocol for ribosome profiling of low input samples"

---

Andreas Meindl[1], Markus Romberger[1], Gerhard Lehmann[2], Norbert Eichner[2], Leon Kleemann[3], Jie Wu[3], Johannes Danner[2], Maria Boesl[1], Mikhail Mesitov[4], Gunter Meister[2], Julian König[4], Sebastian Leidel[3], Jan Medenbach[1]

[1] Regensburg Center for Biochemistry, University of Regensburg, Regensburg, Germany; [2] Biochemistry I, University of Regensburg, Regensburg, Germany; [3] Department of Chemistry, Biochemistry and Pharmaceutical Sciences, University of Bern, Bern, Switzerland; [4] Institute of Molecular Biology (IMB), Mainz, Germany

**Running title**

High sensitivity ribosome profiling

**Keywords**

Ribosome profiling, translation, sequencing library preparation

### Supplemental Figures

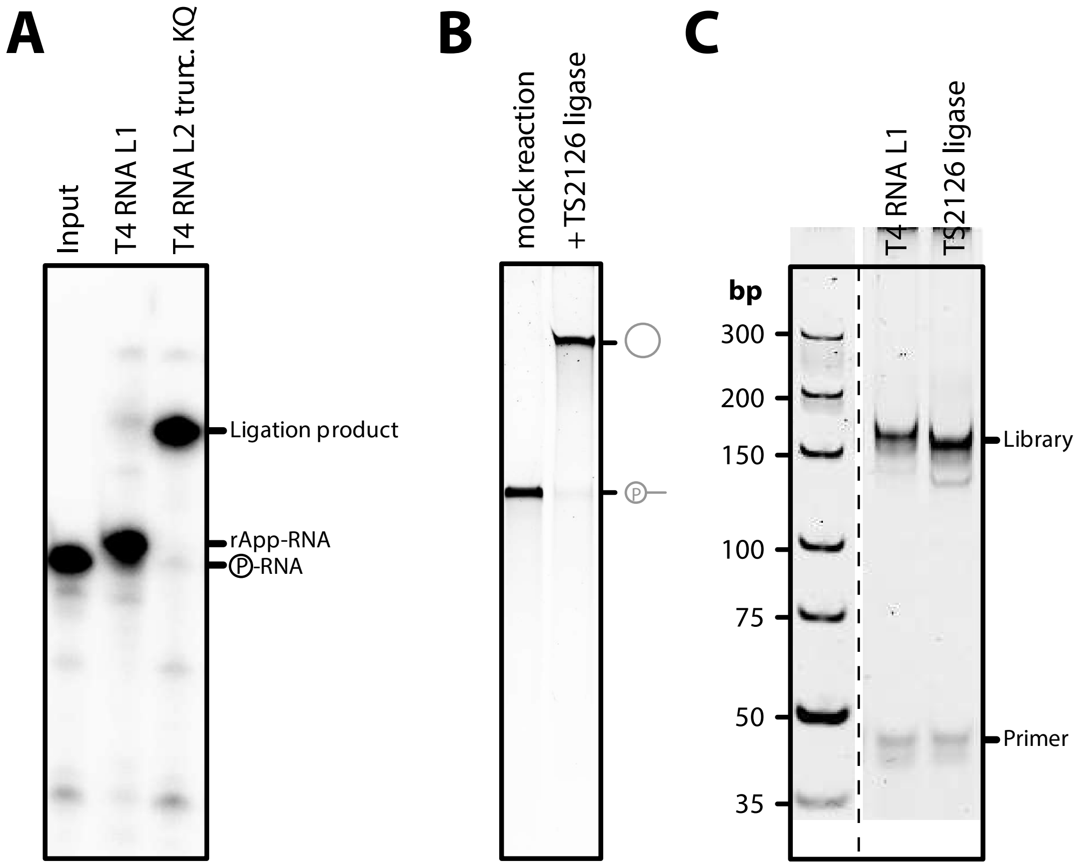


**Supplemental Figure 1 - activity testing of ligases**. **A:** Comparison of T4 RNA ligase 1 (T4RNL1) and T4 RNA ligase 2 (T4RNAL2, aa 1-249, K227Q) activity. 5 pmol of a $^{32}$P radiolabeled RNA (P-RNA, 32 nt) was ligated to 20 pmol of a pre-adenylated DNA oligonucleotide. After incubation over night at 16°C, the reaction products were purified, separated by denaturing PAGE, and visualized by autoradiography. **B:** Circularization of 10 pmol of a 5' phosphorylated 80 nt DNA oligonucleotide by TS2126 ligase. Linear substrate and circular product are schematically indicated on the right. **C:** T4 RNA ligase 1 and TS2126 ligase exhibit comparable performance in sequencing library preparation efficiency. 5 pmol of RPFs were subjected to sequencing library preparation using either T4 ligase 1 for ligation of a second adapter to the cDNA, or Ts2126 ligase for circularization of the cDNA. After 10 cycles of PCR, the final library was separated by PAGE and visualized by ethidiumbromide staining.

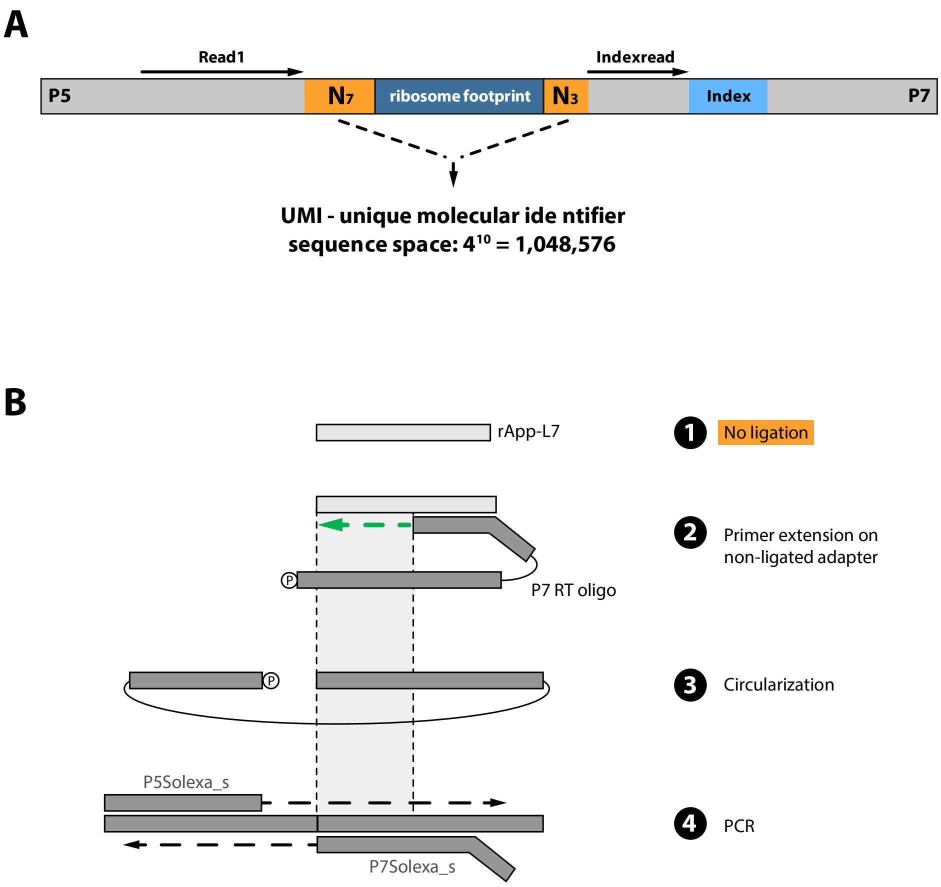


**Supplemental Figure 2 – UMI positions and generation of 'empty' amplicons**. **A:** Schematic depiction of the amplicon generated during library preparation. The ribosome protected fragment (ribosome footprint) in the center of the amplicon is highlighted in blue, the flanking, degenerate sequences used as unique molecular identifiers (UMIs) in orange. Position of the index sequence for experimental multiplexing is highlighted in light blue. The binding sites for primers employed for sequencing are depicted at the top. **B:** Free rApp-L7 adapters (1) can generate empty amplicons during sequencing library preparation (depicted schematically). They serve as a template for the extension of the P7 RT oligo during reverse transcription (2). After circularization (3) this serves as a PCR template, generating products that lack an RPF-derived sequence ('empty' amplicons) (4).

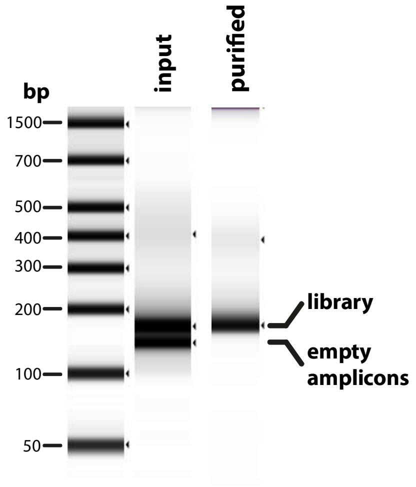


**Supplemental Figure 3 – Gelpurification of the sequencing library**. Tape station analysis of the sequencing library before (input) and after agarose gelpurification (purified) using a Blue Pippin (Sage Science). Empty amplicons and products containing RPF-derived sequence inserts (library) are marked on the right.

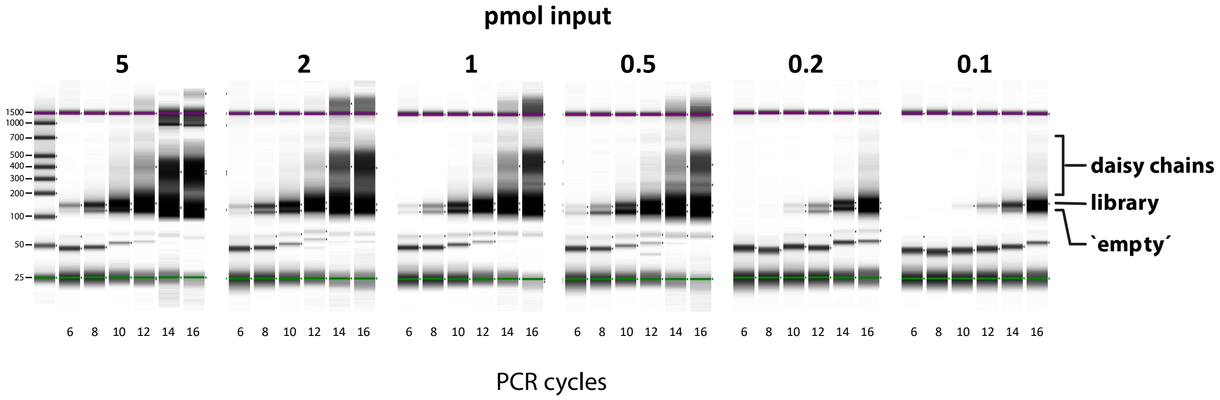


**Supplemental Figure 4 – Sequencing library preparation in non-developer lab**. Tape station analysis products from scouting PCRs of libraries prepared from different amounts of input material (as noted above each gel). Input samples are identical to Figure 3. Provided below each lane are the number of PCR cycles that were performed. The positions of high molecular weight heteroduplex DNA ('daisy chains', derived from over-amplification of the libraries) and 'empty' amplicons are indicated on the right, sizes of the molecular weight marker are indicated on the left.

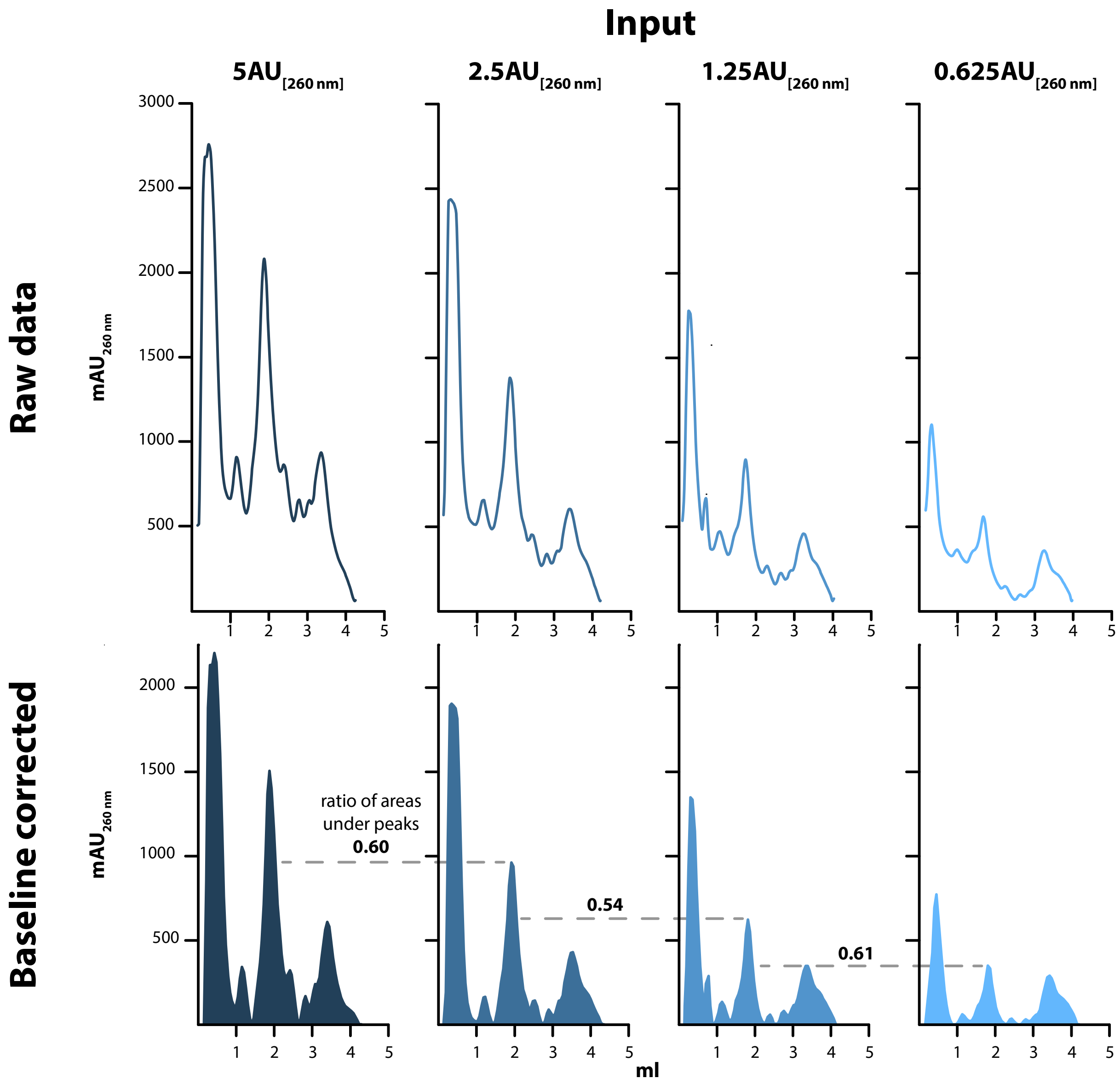


**Supplemental Figure 5 – Baseline correction after sucrose density gradient ultracentrifugation**. Different amounts of K562 extract (serial dilution as indicated above each panel) were separated on linear sucrose gradients. Top: raw UV absorbance traces of the gradients during continuous fractionation, bottom: baseline corrected UV traces.

**Supplemental Table 1: List of primers for sequencing library preparation**. Abbreviations: 3ddC – 3' dideoxy Cytosine, 5Phos – 5' phosphorylated, 5rApp – 5' riboadenylated, * - phosphorothioate bond, iSp19 – internal PEG linker, Index – position of 6 nt index sequence for experimental multiplexing.

| Oligo | Sequence |
|---|---|
| **rApp-L7** | /5rApp/NNNAGATCGGAAGAGCACACGTCTGA/3ddC/ |
| **P7 RT Oligo** | /5Phos/NNNNNNNAGATCGGAAGAGCGTCGTGT/iSp9/GATTCAGACGTGTGC |
| **P5_s** | ACACGACGCTCTTCCGATC*T |
| **P7_s** | GGAGTTCAGACGTGTGCTCTTCCGATC*T |
| **P5** | AATGATACGGCGACCACCGAGATCTACACTCTTTCCCTACACGACGCTCTTCCGATCT |
| **TrueSeq_P7_Index** | CAAGCAGAAGACGGCATACGAGAT-Index-GTGACTGGAGTTCAGACGTGTGCTCTTCC |

**Supplemental Table 2: Biotinylated rRNA depletion oligos.** HPLC purified oligos were ordered from IDT containing 5' Biotin TEG (15 atom triethylene glycol spacer).

| Oligo | Sequence |
|---|---|
| **1** | /5BiotinTEG/TG CGC CGC GAC CGG CTC CGG GAC GGC TGG |
| **2** | /5BiotinTEG/CA CTC GCC GAA TCC CGG GGC CGA GGG AGC |
| **3** | /5BiotinTEG/TG GGG GGC CCA AGT CCT TCT GAT CGA GGC |
| **4** | /5BiotinTEG/CC CGG GGC TAC GCC TGT CTG AGC GTC GCT |
| **5** | /5BiotinTEG/CC GCC TGG GAA TAC CGG GTG CTG TAG GCT |
| **6** | /5BiotinTEG/CG CTG CGA TCT ATT GAA AGT CAG CCC TCG |
| **7** | /5BiotinTEG/CG CCG AGG GCG CAC CAC CGG CCC GTC TCG |
| **8** | /5BiotinTEG/CG CCG GGT TAA GGC GCC CGA TGC CGA CGC |
| **9** | /5BiotinTEG/TC GAA TAC AGA CCG TGA AAG CGG GGC CTC |
| **10** | /5BiotinTEG/TA CGG TGG CCA TGG AAG TCG GAA TCC GCT |
| **11** | /5BiotinTEG/TG CAG TTA AAA AGC TCG TAG TTG GAT CTT |
| **12** | /5BiotinTEG/TT TCA CTG ACC CGG TGA GGC GG |
| **13** | /5BiotinTEG/TC GCT TCT GGC GCC AAG CGC CC |
| **14** | /5BiotinTEG/GG TTG GCC TCG GAT AGC CGG TC |
| **15** | /5BiotinTEG/TC GGC CGA GGT GGG ATC CCG AG |
| **16** | /5BiotinTEG/TA AAC GAT GCC GAC CGG CGA TGC GGC GGC |
| **17** | /5BiotinTEG/TC CGC CAC GCA GTT TTA TCC GGT AAA GCG |
| **18** | /5BiotinTEG/TC GGC GAC GAC CCA TTC GAA CGT CTG CCC |
| **19** | /5BiotinTEG/AC AAA GCA CCC AAC TTA CAC TTA GGA GAT |
